# CellWell: A micropatterned biphasic nanocomposite platform for culturing chondrocytes

**DOI:** 10.1101/790030

**Authors:** Ram Saraswat, Ishara Ratnayake, E. Celeste Perez, William M. Schutz, Zhengtao Zhu, S. Phillip Ahrenkiel, Scott T. Wood

## Abstract

**Graphical Abstract:** 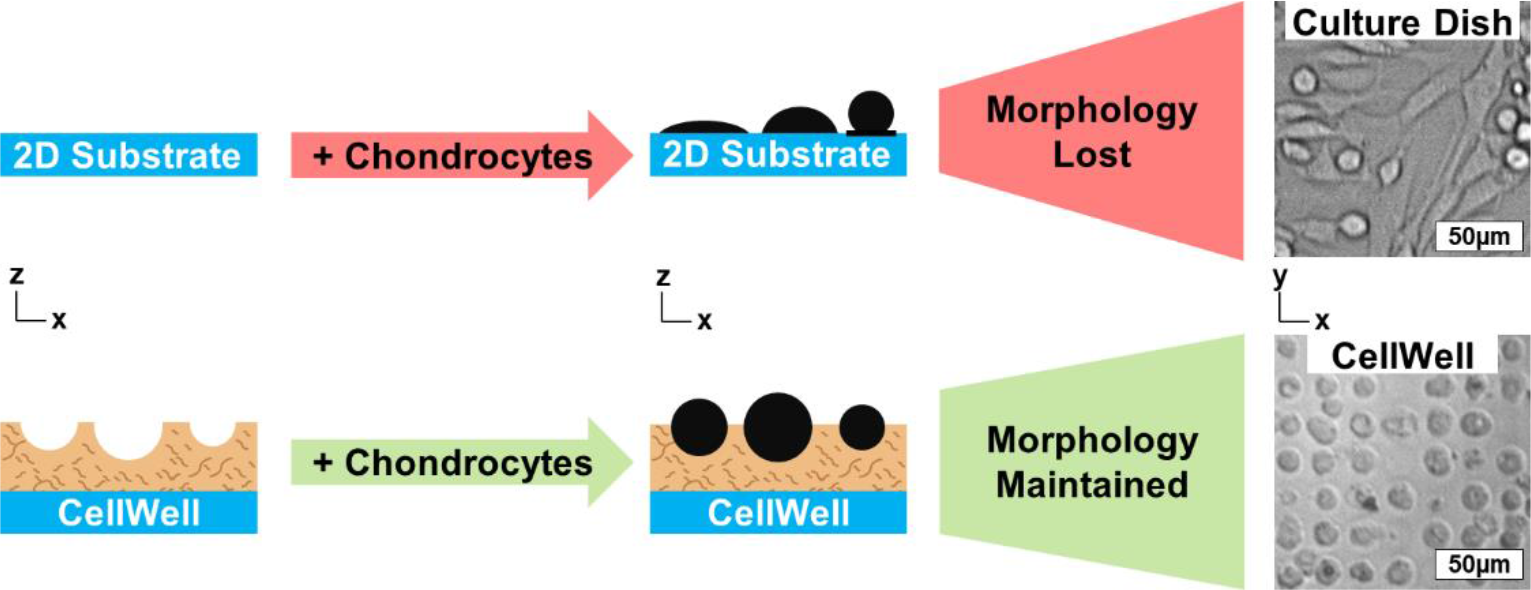

**Abstract:** We present a unique micropatterned nanocomposite cell culture platform to model articular cartilage that is suitable for high-throughput single-cell analyses using standard imaging techniques. This platform, the CellWell, is constructed out of a thin, optically transparent substrate that is lithographically micropatterned with a network of wells sized to fit individual cells. The substrate material consists of a thin layer of agarose hydrogel embedded with polyvinyl alcohol nanofibers. The geometries of the wells are designed to reinforce a physiological morphology, thereby combining the physiological advantages of 3D culture systems with the practical advantages of 2D systems. CellWells were found to have compressive moduli of 144 ± 11.5 kPa and 158 ± 0.6 kPa at strain rates of 5 μm/s and 15 μm/s. The compressive moduli were determined at two different strain rates to allow for comparison of CellWell stiffness with published values of pericellular matrix and with observed values of articular cartilage, which could not be indented at the same rate. Articular chondrocytes seeded in a CellWell were found to maintain their spheroidal morphology more effectively than those seeded in monolayer cultures and to be more easily imaged than those seeded in a 3D scaffold of identical thickness. Through its ease of use and ability to maintain the physiological morphology of chondrocytes, we expect that the CellWell will enhance the clinical translatability of future studies conducted using this culture platform.

## 1. Introduction

Osteoarthritis (OA) is a painful disease of the articular joints that is primarily characterized by the degradation of the extracellular matrix (ECM) in the articular cartilage [1]. To date, surgical restoration techniques used for cartilage repair do not regenerate hyaline articular cartilage. Although symptoms can improve temporarily after surgical repair, 85% of patients progress to failure within 7.5 years or less [2]. There are currently no known medical treatments that effectively address the underlying molecular causes of OA. Current pharmaceutical treatment options are limited to the use of analgesics like non-steroidal anti-inflammatory drugs (NSAID) and intra-articular corticosteroid injections to reduce the pain associated with inflammation which only provide temporary relief and can have negative consequences with long-term use [3–9]. Articular chondrocytes are the only cells in the articular cartilage and are responsible for the maintenance of cartilage homeostasis between digestion and replacement of old or damaged tissue components. It is well-accepted that a loss of this homeostatic balance is responsible for the development of OA [10].

Animals models have long been the gold standard for understanding the progression of osteoarthritis. However, they are also associated with concerns of ethical issues regarding the treatment of animals, cost and management issues, anatomical differences of cartilage in animals compared to humans, and age variations of animal species at the time of testing [11–25].

Due to the problems associated with animal models, chondrocytes have been studied *in vitro* using either standard two-dimensional (2D) or any number of three-dimensional (3D) cell culture techniques. When plated on standard 2D platforms, chondrocytes tend to rapidly lose their canonical spheroidal morphology due to the adherent chondrocyte cells being on a hard and flat substrate, and adopt a fibroblastic phenotype within 14 days of culture [26–29]. These morphological changes can result in substantial changes to chondrocyte architecture, including the length, density, and distribution of cortical actin fibers. The *in situ* chondrocyte shown in Fig. 1(A) can be seen to have an actin network that is more similar to the chondrocyte displaying the canonical phenotype in Fig. 1(B), than to that of the chondrocyte in Fig. 1(C) which has a spread morphology that is more typical of chondrocytes in standard monolayer culture. Note that the *in situ* chondrocyte in Fig. 1(A) is not flat, but that the turbidity of the tissue prevented imaging its full thickness, thereby illustrating the difficulties inherent to imaging cells in 3D samples. It has long been known that forcing chondrocytes to adopt a rounded morphology leads to enhancement of a chondrocytic phenotype *in vitro*; however, the techniques utilized previously have all relied upon restriction of binding area on a 2D substrate to prevent spreading rather than active promotion of a rounded phenotype in a way that does not inherently limit adhesion [30, 31].

**Fig. 1:**
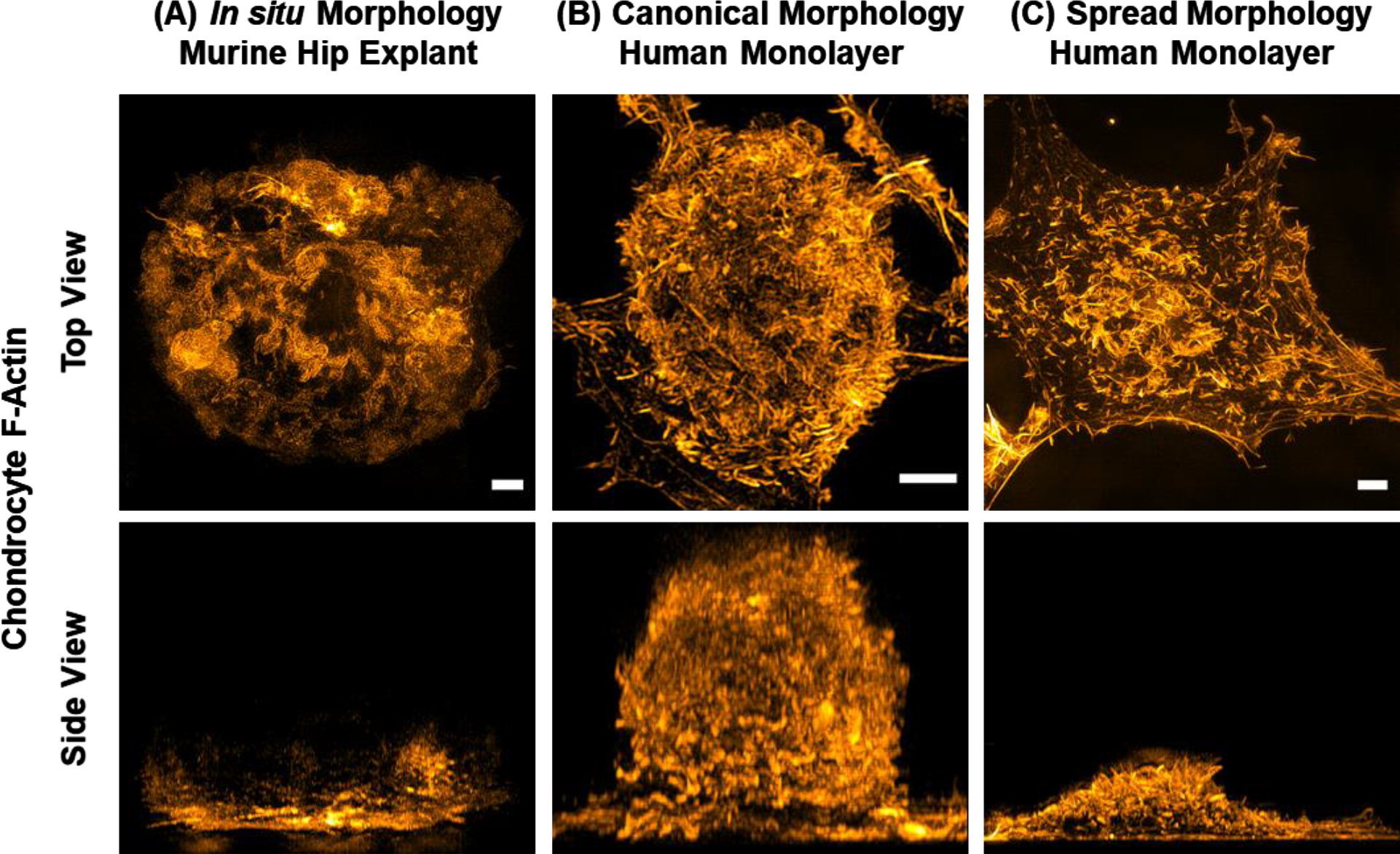
Chondrocyte morphology influences internal architecture. Murine femoral cap hip explants (A) and primary human articular chondrocytes (B) and (C) were plated on fibronectin-coated coverglass using standard 2D cell culture techniques, fixed with 4% paraformaldehyde, stained with ActinRed 555 Ready probes® Reagent (ThermoFisher) and imaged in super-resolution using 3D structured illumination microscopy (3D-SIM) on a GE DeltaScan™ OMX SR microscope. Maximum intensity projections of volumetric image stacks are shown. Note in the side view of (A) that the turbidity of the tissue prevented imaging of all but the surface-most structures in the explant images, a common problem in imaging of both natural tissues and 3D cell cultures. Scale bars are 2 μm and apply to both top and side projections.

In this study, we establish the proof of concept for a unique micropatterned nanocomposite cell culture platform, the CellWell, which consists of a thin film micropatterned nanofiber-embedded hydrogel substrate that fits a single cell within each well and facilitates high throughput fluorescence imaging of chondrocytes. The substrate composition was chosen to recapitulate the ECM of articular cartilage wherein a hydrogel models cartilage proteoglycans and embedded nanofibers model collagen II fibers. Our goals for the design of the CellWell included: (1) designing the wells such that their geometries reinforce the canonical spheroidal chondrocyte morphology for each cell; (2) matching the mechanical stiffness of articular cartilage ECM or the chondrocyte pericellular matrix (PCM) as closely as possible; (3) matching the diameters of the embedded nanofiber diameters as closely as possible to those of the native collagen II fibers; and (4) ensuring compatibility with traditional cell culture and live-cell imaging techniques.

## 2. Materials and Methods

### 2.1 Materials Required

All the reagents for hydrogels and fibers were purchased from Fisher Scientific, USA unless otherwise noted. All the reagents for Transmission Electron Microscope (TEM) sample preparation were purchased from Ted Pella, Inc. (Redding, CA). Normal human articular cartilage samples from the ankle joint were obtained from the Thurston Arthritis Research Center (TARC) at the University of North Carolina (UNC) School of Medicine through the Department of Biochemistry at Rush University Medical Center (Chicago, IL) from the Gift of Hope Organ and Tissue Donor Network (Elmhurst, IL), and from Dakota Lions Sight and Health (Sioux Falls, SD).

### 2.2 Hydrogel Preparation

Agarose hydrogels (5% w/v) were prepared with slight modifications to the method described by H.M. Pauly et al. [32]. Poly(vinyl alcohol) (PVA) (15% w/v) hydrogels were prepared, based upon the method described by S. Jiang et al. [33].

### 2.3 Electrospun Nanofiber Preparation

PVA solution was prepared based upon the method described by A.G. Destaye et al. [34], and the electrospun nanofibers were obtained using the setup described by S.Mishra et al.[35]. The electrospun nanofibers were then crosslinked under via glutaraldehyde vapors for 48 hrs [34] in a vacuum desiccator.

### 2.4 CellWell Design and Manufacturing

The diameter of 8,375 chondrocytes from 18 independent donors was measured using a Countless II FL cell counter (ThermoFisher). This information was used to establish the well diameters of the CellWell. Computer-aided design (CAD) files were generated in SolidWorks consisting of an array of circles of 12, 15, and 18 μm diameters and used to generate the photomask pattern, as shown in Fig. 2. The design was chosen such that the distance between any two consecutive wells varied from 2 μm to 15 μm.

**Fig. 2:**
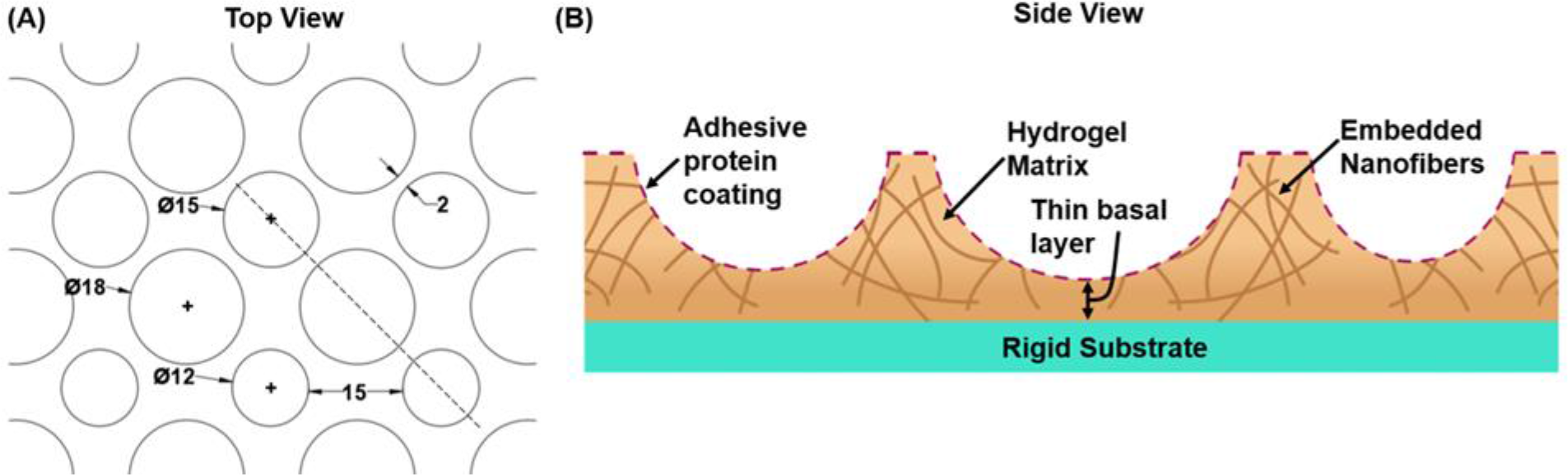
CellWell design schematics. (A) Representative section of photomask design. Units are in μm. (B) Cross-sectional representation of CellWell along the dashed line in (A).

Micropatterned silicon wafers were obtained from the Utah Nanofab core lab at the University of Utah, USA, and standard contact lithography techniques were utilized to generate PDMS CellWell stamps [36]. PDMS stamps were sterilized in an autoclave at 121 °C for 23 minutes [37]. “Containment chambers” were microfabricated with 15 μm-tall walls, in which the CellWell casting process took place. These walls were thus constructed to be ~8 μm taller than the hemispheroids in the stamps to provide room for several microns of material to separate the basal surface of the cells from the underlying cover glass without adding excessive bulk that can confound imaging experiments conducted on standard inverted microscopes. To cast CellWells, molten agarose solution, mixed with finely chopped crosslinked PVA nanofibers, was poured into a containment chamber, and the composite molten solution was stamped with a PDMS stamp at 4 °C for 6 mins. The stamp was then removed, revealing the bare CellWell. CellWells were then immediately hydrated with PBS-1x solution, UV sterilized for 30 mins, and coated with 10 μg/ml each of purified human plasma fibronectin and human placenta collagen type VI (Rockland Immunochemical) for 30 minutes at 37 °C.

### 2.5 Cell Culture

Articular cartilage donors (N=4) had an age range of 42-77, a male/female ratio of 2/2, Collins scores ranging from 0-2, and no known history of OA. Primary human articular chondrocyte isolation from de-identified ankle articular cartilage was performed as described previously [38]. Briefly, chondrocytes were isolated using sequential digestion with Pronase and collagenase, then plated in 35 mm tissue culture dishes and pre-incubated for 2 days to allow the cells to recover from the digestion process. Full-thickness articular cartilage explants were prepared before enzymatic digestion of the tissue using a 5 mm biopsy punch. Chondrocytes were gently lifted from the substrate using a 1 hr treatment with Pronase and collagenase and then seeded onto CellWells or control substrates. Chondrocytes were plated on top of tissue culture polystyrene or 15 μm-thick non-patterned agarose for 2D controls and encapsulated within thickness-matched agarose for 3D control samples [39]. In all cases, chondrocytes were seeded with a density of 2×10^5^ cells/cm^2^. Culture media was replaced at one hour after initial seeding, after which cells were incubated continuously for 23 hrs before imaging on an Olympus IX71 inverted epifluorescence microscope with a 20X, 0.46 N.A. objective (Olympus) and an Andor iXon Ultra EMCCD camera (Andor USA). Viability of chondrocytes was assessed in CellWells at 24 hrs post-seeding using a fluorescent ReadyProbes^®^ Cell Viability Imaging Kit (ThermoFisher).

### 2.6 Mechanical Characterization

The viscoelastic properties of CellWells and articular cartilage were analyzed using an Asylum Research MFP 3D Atomic Force Microscope (AFM), with Igor Pro v6.37. Borosilicate glass spheres (4.8 ± 0.3 μm diameter; SPI Supplies) were attached to the tip of AFM cantilevers (force constant in the range 0.04-0.7 N/m; All-In-One-Al-Tipless, Budget Sensors) using epoxy, and spring constants for each cantilever were determined thermally [40, 41] before experimentation. Stress relaxation of Agarose CellWells (N=3) and articular cartilage explants (N=3) was performed using a 5 μm/s approach velocity and 60 s relaxation time, as depicted in Fig. 3. The indentation phase was utilized for all elastic parameter calculations, while the relaxation phase was utilized for all viscosity parameter calculations.

**Fig. 3:**
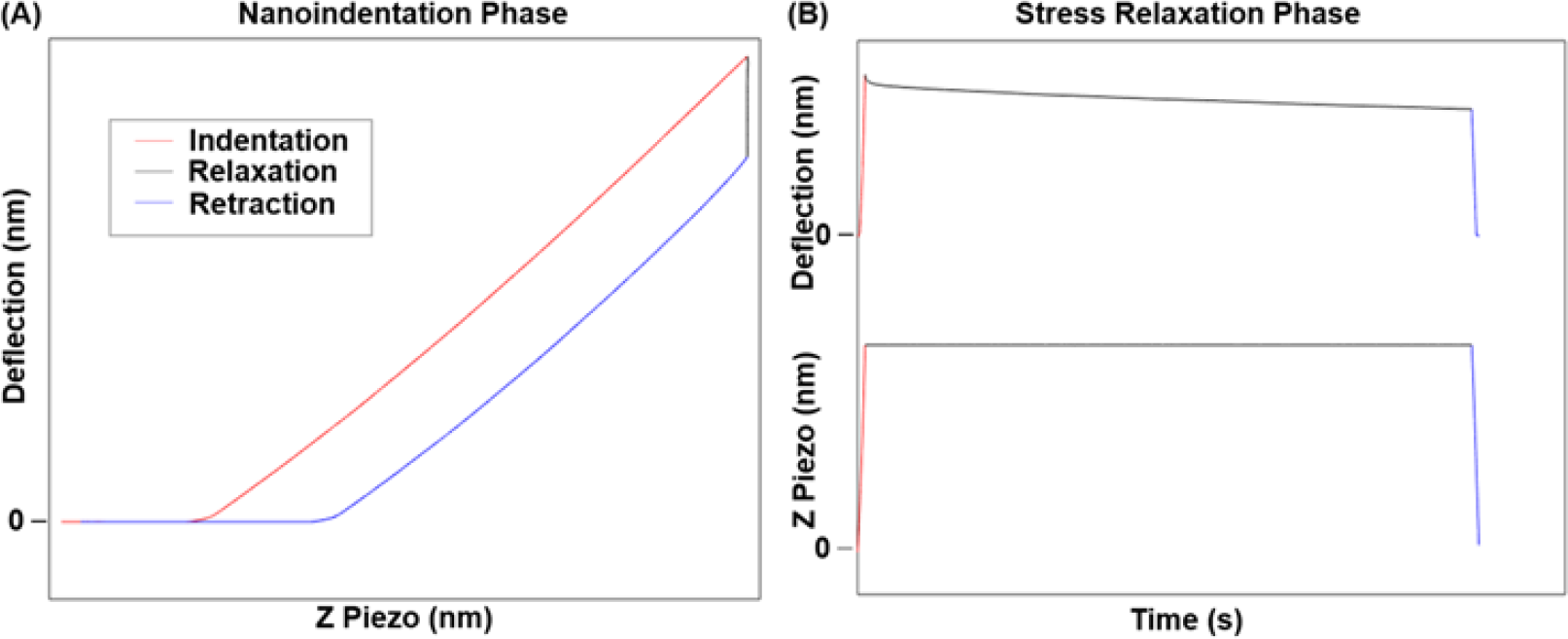
AFM Experimental Design Schematics. (A) Nanoindentation Phase (B) Stress Relaxation Phase

Once the raw curves were obtained, the raw deflection curves were converted to force curves using Hooke’s Law;

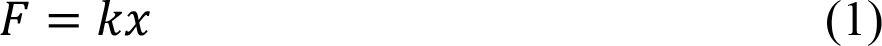

Where *F* is the force, *x* is the deflection of the cantilever, and *k* is the cantilever spring constant determined thermally.

To analyze the viscoelastic properties, a modified version of the Standard Linear Solid (SLS) Model as described by E.M. Darling et al. [42] was used.

All the force fittings were done as per the method described by E.M. Darling et al. [42], described by the following equations:

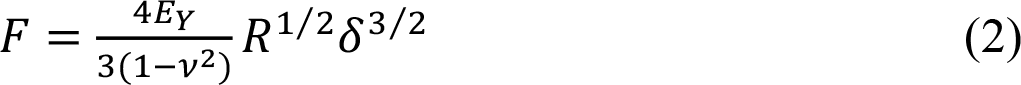

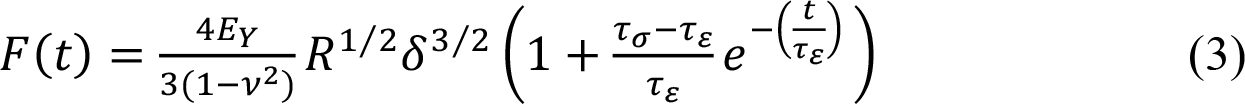

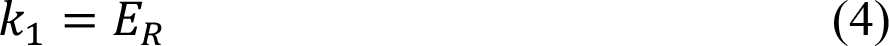

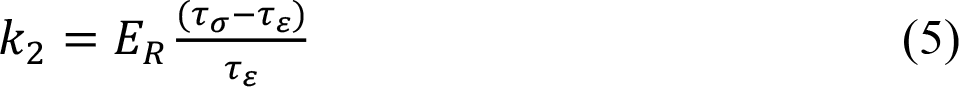

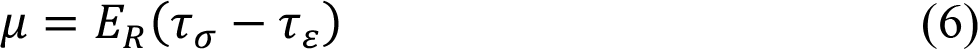

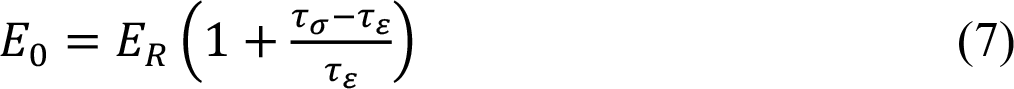

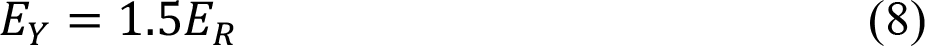

Where *E*_*Y*_ is the Hertz Compressive Moduli, *F* is the applied force during indentation, *v* is the Poisson’s ratio, *R* is the radius of the indenter (2.5 μm), *F*(*t*) is the force measured as a function of time during stress relaxation, *E*_*R*_ is the SLS Relaxation Moduli, *τ*_*σ*_ is the relaxation time under constant load, *τ*_*∊*_ is the relaxation time under constant deformation, *k*_1_ and *k*_2_ are the Kelvin spring elements, *μ* is the apparent viscosity and *E*_0_ is the instantaneous moduli.

As per E.M. Darling et al. [42], Eq (2) fits the Hertz equation, and Eq (3) fits the Standard Linear Solid (SLS) Model. The Poisson’s ratio of agarose and cartilage were both assumed to be 0.33 [43], and calculations were performed based on measurements at 1,500 nm of indentation depth (10% compressive strain for CellWells).

Agarose CellWells (N=3) were separately indented at 15 μm/s to allow for direct comparison of compressive moduli with that of human pericellular matrix published by E.M. Darling et al. [44]. For this comparison, the compressive moduli were obtained using Eq (2) at 8% compressive strain.

### 2.7 Optical Transmittance

Optical transmittance of agarose, PVA nanofibers, and nanofiber-embedded CellWells (N=3 each) in the visible range was measured using a Video Spectral Comparator (VSC). Transmittance values were normalized against coverglass controls.

### 2.8 Nanofiber Characterization

PVA and ankle cartilage collagen II nanofibers were imaged using a JEOL JEM-2100 LaB6 transmission electron microscope (TEM), and diameters were measured using FIJI ImageJ v1.52n. PVA nanofibers were prepared as described in 2.3 and mounted on TEM grids for imaging. For collagen II diameter measurements, articular cartilage explants from the ankle were fixed with 2% PFA and 2.5% glutaraldehyde in 0.1 M cacodylate solution for 1 hr, followed by rinsing with sodium cacodylate buffer (0.1 M, pH 7.2) three times 5 min each. Then the tissues were postfixed with 0.5% OsO_4_ and 0.5% potassium ferrocyanide for 30 min. After rinsing with cacodylate buffer, the tissues were dehydrated in a series of ethanol solutions (50%, 70%, 90% and 100% for 20 min each). The tissues were infiltrated with a mixture of ethanol and Araldite (2:1, 1:1, 1:2 ratios for 2 hrs each) and cured with a fresh Araldite resin at 60°C for 48 hr. Sections of 70 nm thickness were cut with an ultramicrotome (RMC Powertome XL), mounted on TEM grids, and stained with uranyl acetate and lead citrate.

### 2.9 Statistical Analysis

Data from a minimum of three independent experiments were analyzed for all statistical analyses. Linear regression was used to determine the quality of fit of Hertzian and viscoelastic models to AFM indentation data. Cell morphology measurements were collected on a single-cell basis. A Shapiro-Wilk test was used to determine whether data sets had normal or lognormal distributions. A Kruskal-Wallis test with Dunn’s multiple comparisons post-hoc test was used to determine effects since data sets were found to have neither normal nor lognormal distributions. All statistical analyses were performed in GraphPad Prism v8.2.1.

## 3. Results

### 3.1 Chondrocyte Diameter

As shown in Fig. 4(A), human chondrocytes were found to have a mean diameter of 14.6μm ± 2.1 μm (S.D.), and this data represented the diameter measurements from 8,375 cells over n=18 individual donors. The chondrocyte diameter distribution of average donors depicted a Full Width at Half Maxima (FWHM) of 12-18 μm, as shown in Fig. 4(B). These measurements served as the basis for the selection of CellWell diameters of 12, 15, and 18 μm.

**Fig. 4:**
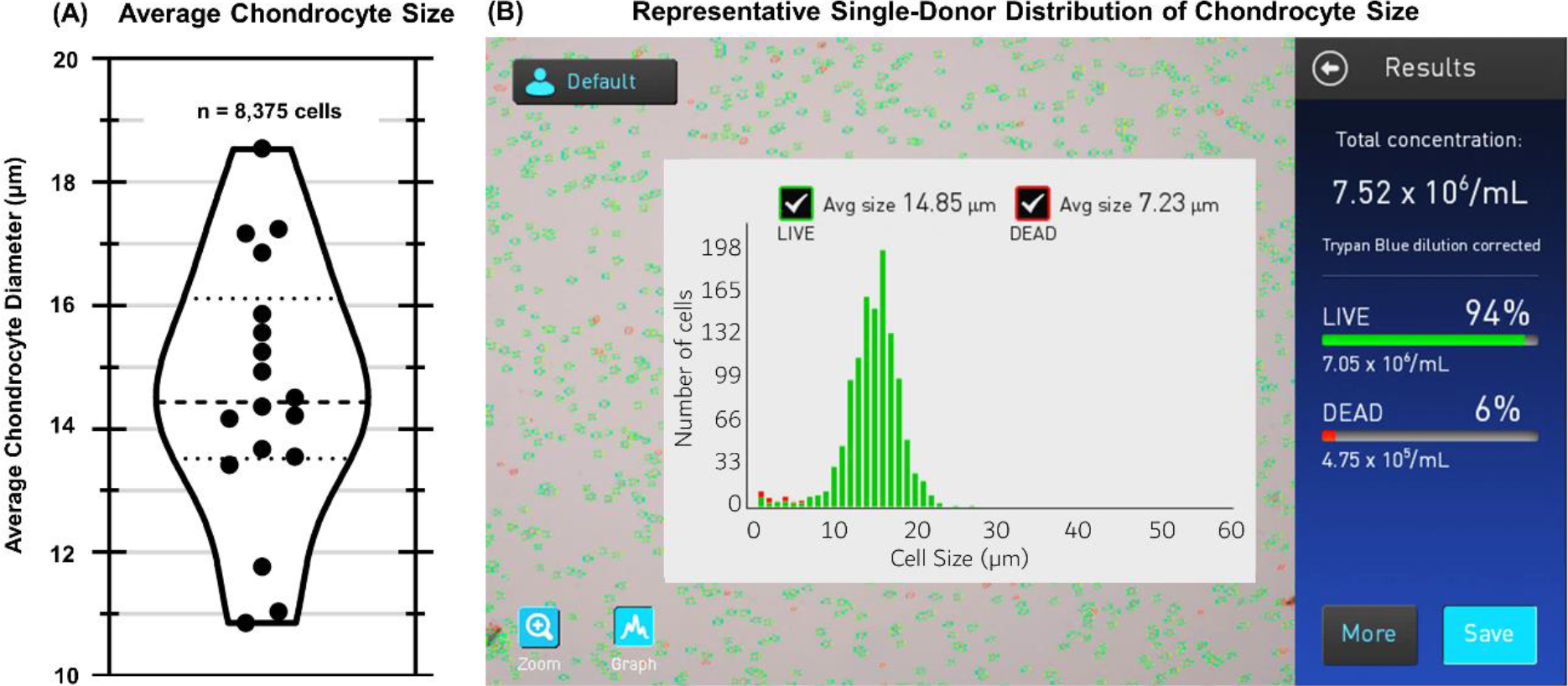
Chondrocyte diameter distribution. (A) Primary human articular chondrocytes were found to have a mean diameter of 14.6±2.1 μm (S.D.) in suspension, with average diameters on a per-donor basis ranging from ~11-19 μm. (B) Cell counter screenshot screen showing a representative single-donor chondrocyte diameter distribution.

### 3.2 CellWell Manufacturing

A Keyence VK-X250 optical profilometer was used to measure the dimensions of CellWell features (N=10) . One of the limitations of our profilometer was that it could only work with dry samples and so we expected shrinkage effects in our CellWells due to the fact that the gelation mechanism of agarose is solely based on the physical hydrogen-bond networks [45–48]. Thus, to ensure the fidelity of collected data, CellWells for these measurements were made out of PVA because PVA was made by freeze-thaw method as described in 2.3, and frozen samples were able to be utilized to minimize the loss of feature height due to hydrogel drying compared to CellWells made of agarose. Fig. 5(A) shows scanning electron microscopy (SEM) images of the lithographic patterns on silicon that were used to create CellWell stamps. Fig. 5.(B) shows a phase contrast image of a CellWell with three sizes of wells precisely sized to fit individual articular chondrocytes and Fig. 5.(C) shows the cross-sectional profiles of individual wells as measured by optical profilometry. Although frozen PVA CellWells were observed to have a decrease in well height by 7-15% due to imaging in the dried state, it can be easily seen in Fig. 5(C) that the geometry of the wells is hemispheroidal.

**Fig. 5:**
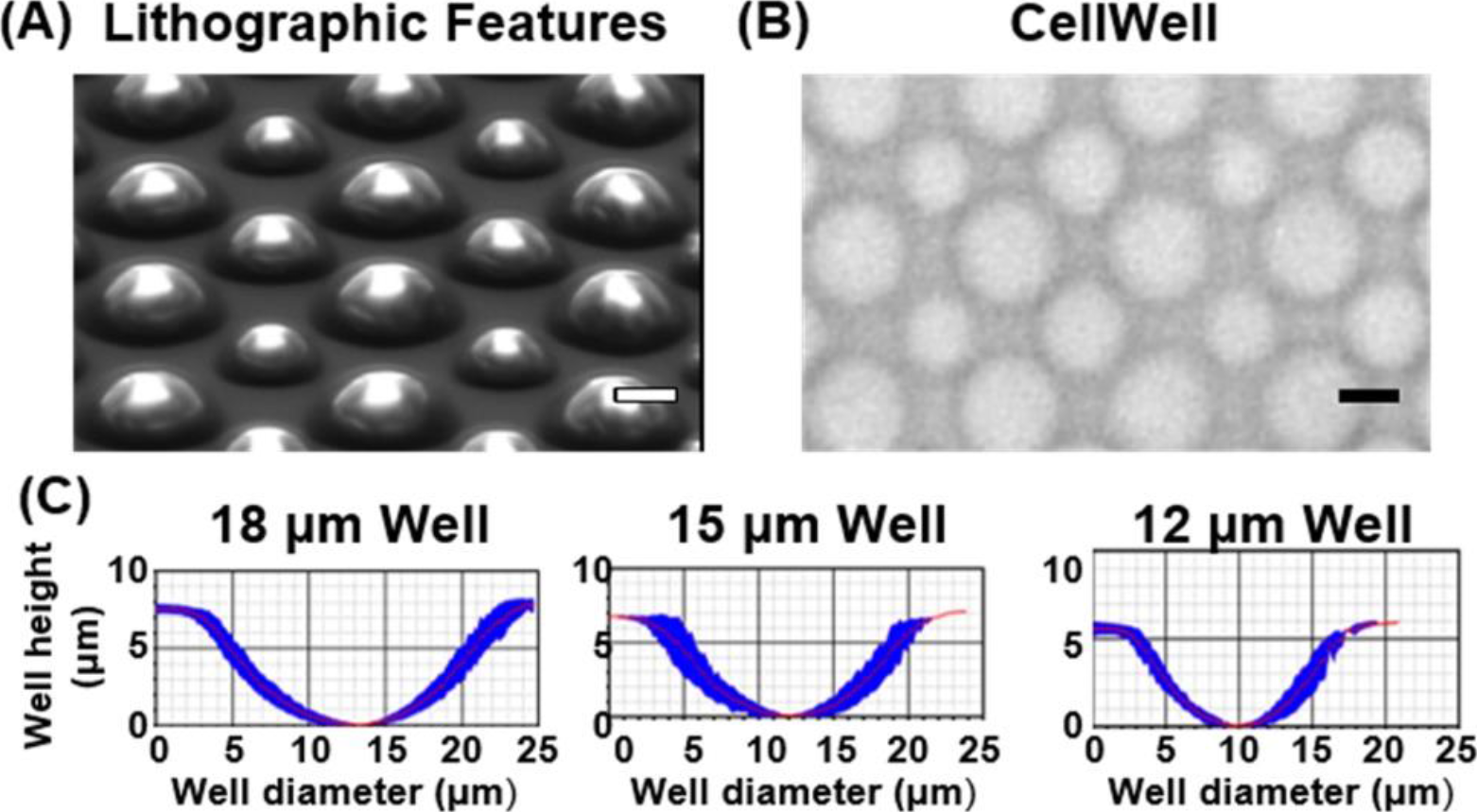
Micropatterned wells match chondrocyte diameters. (A) Scanning Electron Microscopy (SEM) image of lithographic patterns used to create CellWells (B) Phase Contrast Image of CellWell with three sizes of wells precisely sized to fit individual articular chondrocytes; Scale bars10 μm. (C) Optical profilometry cross-sectional profiles of individual wells (mean ± S.D. of n=10 wells each).

### 3.3 Mechanical Characterization

Fig. 6 (A) and (C) depict average indentation curves at 5 μm/s for agarose and cartilage (N=3), respectively, along with both Hertzian (Eq (2)) and viscoelastic SLS model (Eq (3)) fits. Fig. 6 (B) depicts average stress relaxation curves for agarose and cartilage along with the respective SLS fit for each based on Eq (3). These models were used to analyze the mechanical properties of agarose CellWells and ankle articular cartilage shown in Table 1.

**Fig. 6:**
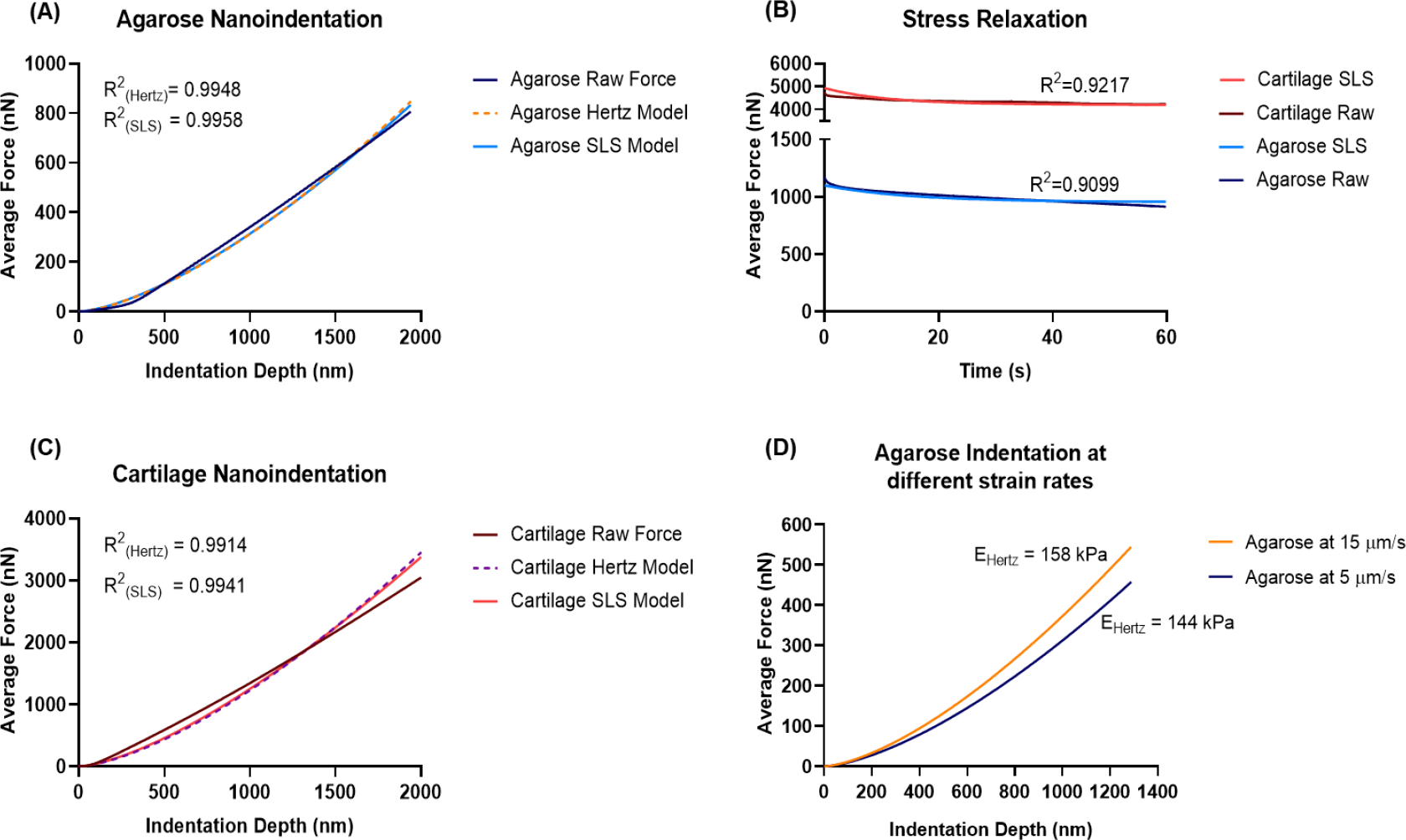
Mechanical characterization of CellWell agarose and ankle articular cartilage. (A) Average Nanoindentation Curves of Agarose CellWells (B) Average Stress Relaxation of Agarose CellWells and Articular Cartilage (C) Average Nanoindentation Curves of Articular Cartilage (D) Average Nanoindentation on Agarose CellWells at different strain rates.

**Table 1.** Comparison of mechanical properties of the Agarose CellWells and articular cartilage (Mean ± S.D.)

| Mechanical Parameter | CellWell | Articular Cartilage |
| --- | --- | --- |
| Hertz Elastic Modulus, $E_Y$ (kPa) | $144 \pm 11.5$ | $488 \pm 102.5$ |
| Relaxation Modulus, $E_R$ (kPa) | $95.8 \pm 7.65$ | $325 \pm 68.3$ |
| Instantaneous Modulus, $E_0$ (kPa) | $175 \pm 24.5$ | $575 \pm 126.5$ |
| $\tau_\sigma$ (s) | $17.3 \pm 1.04$ | $15.0 \pm 4.80$ |
| $\tau_\epsilon$ (s) | $14.3 \pm 0.86$ | $12.8 \pm 4.09$ |
| $k_1$ (kPa) | $95.8 \pm 7.65$ | $325 \pm 68.3$ |
| $k_2$ (kPa) | $20.8 \pm 7.90$ | $58.0 \pm 16.8$ |
| $\mu$ (kPa·s) | $296 \pm 103.6$ | $677 \pm 60.9$ |

Figure 6 and Table 1 shows comparisons of the CellWell with human ankle articular cartilage. While slightly less stiff than cartilage as measured by AFM, the CellWell provides a much more comparable mechanical environment to articular cartilage (70% lower) than commonly used tissue culture polystyrene (10^4^ higher), coverglass (10^5^ higher), or softer hydrogels (10^2^ lower). Thus, the CellWell provides a much more appropriate mechanical environment for the cells than standard monolayer cultures.

To compare the compressive mechanical properties of CellWells with that of human extracellular and pericellular matrices published by E.M. Darling et al. [44], we indented the agarose CellWells at 15 μm/s. As a comparison, the agarose at 5 μm/s was also plotted to depict the effect of strain rate, as shown in Fig. 6 (D). Importantly, the CellWell elastic modulus of 144 ± 12 kPa (S.D.) is very close to the reported 162 ± 22 kPa (S.D.) stiffness of knee cartilage PCM, indicating that it provides a highly appropriate mechanical environment for chondrocytes.

### 3.4 Optical Characterization

In general, the turbidity of 3D samples makes it difficult to image them beyond their surface level. To ensure that the CellWells are optically transparent enough to facilitate clear imaging on an inverted microscope with standard live-cell imaging techniques, we measured the optical transmittance of the CellWells across the visible range (Fig. 7). Even though the nanofibers had a transmittance of only about 50%, the transmittance of the hydrogels was found to be higher than 85%, and the nanofiber embedded hydrogels ranged from 70-85%.

**Fig. 7.**
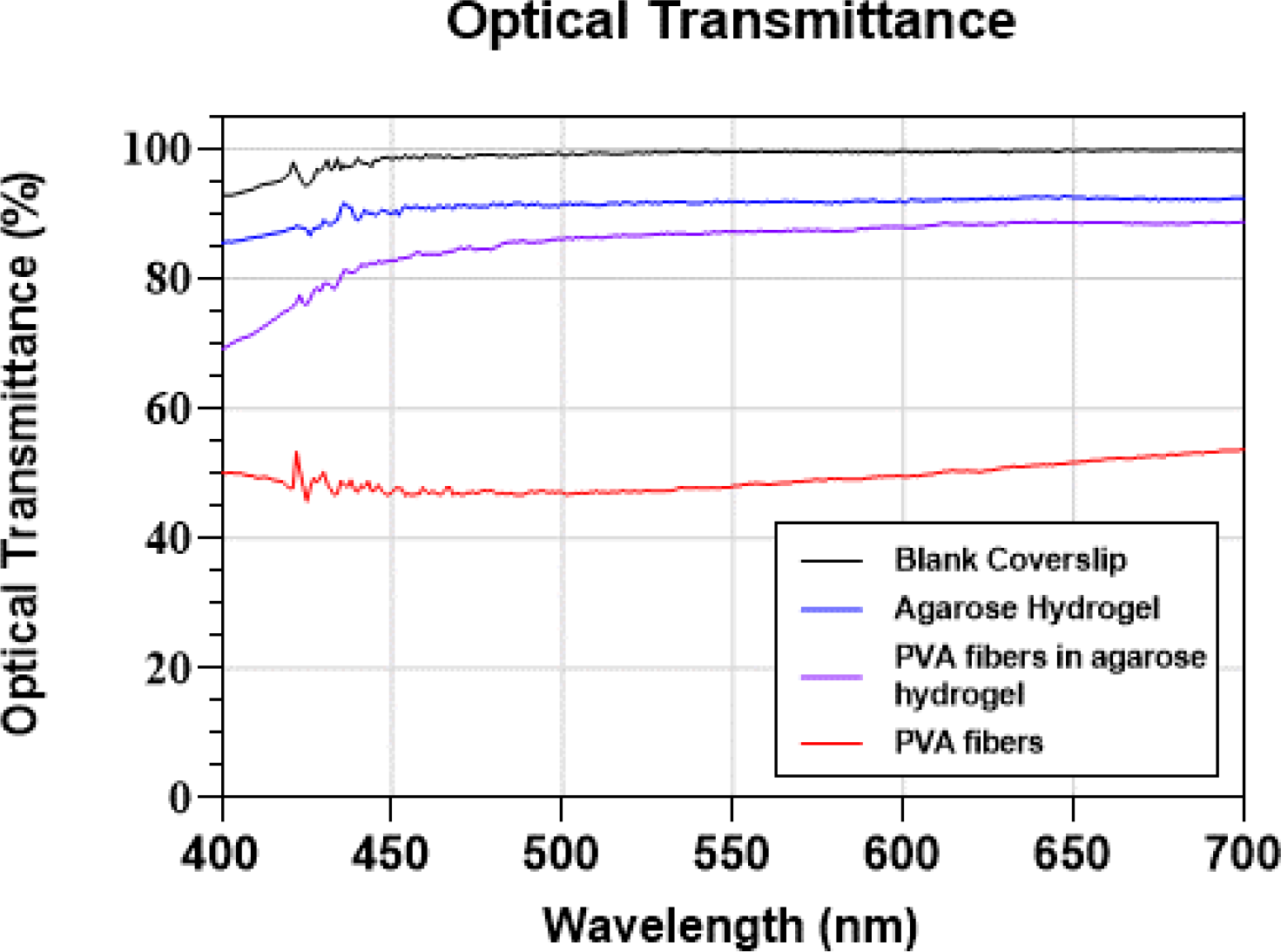
Optical transmittance of the CellWells measured using Video Spectral Comparator, confirming the optical transparency of the CellWells.

### 3.5 Protein Coating

To confirm the adsorption of PCM proteins onto agarose CellWells, Fourier transform infrared spectra (FTIR) of coated CellWells were obtained for samples coated with either fibronectin (FN) or type-VI collagen (Col-VI). The FTIR spectra of an uncoated agarose CellWell and pure PCM proteins were also analyzed and used as controls. As depicted in Fig. 8, the FTIR spectra of agarose CellWells coated with FN or Col-VI was representative of the pure proteins, thus confirming that the PCM proteins were successfully adsorbed onto CellWells.

**Fig. 8.**
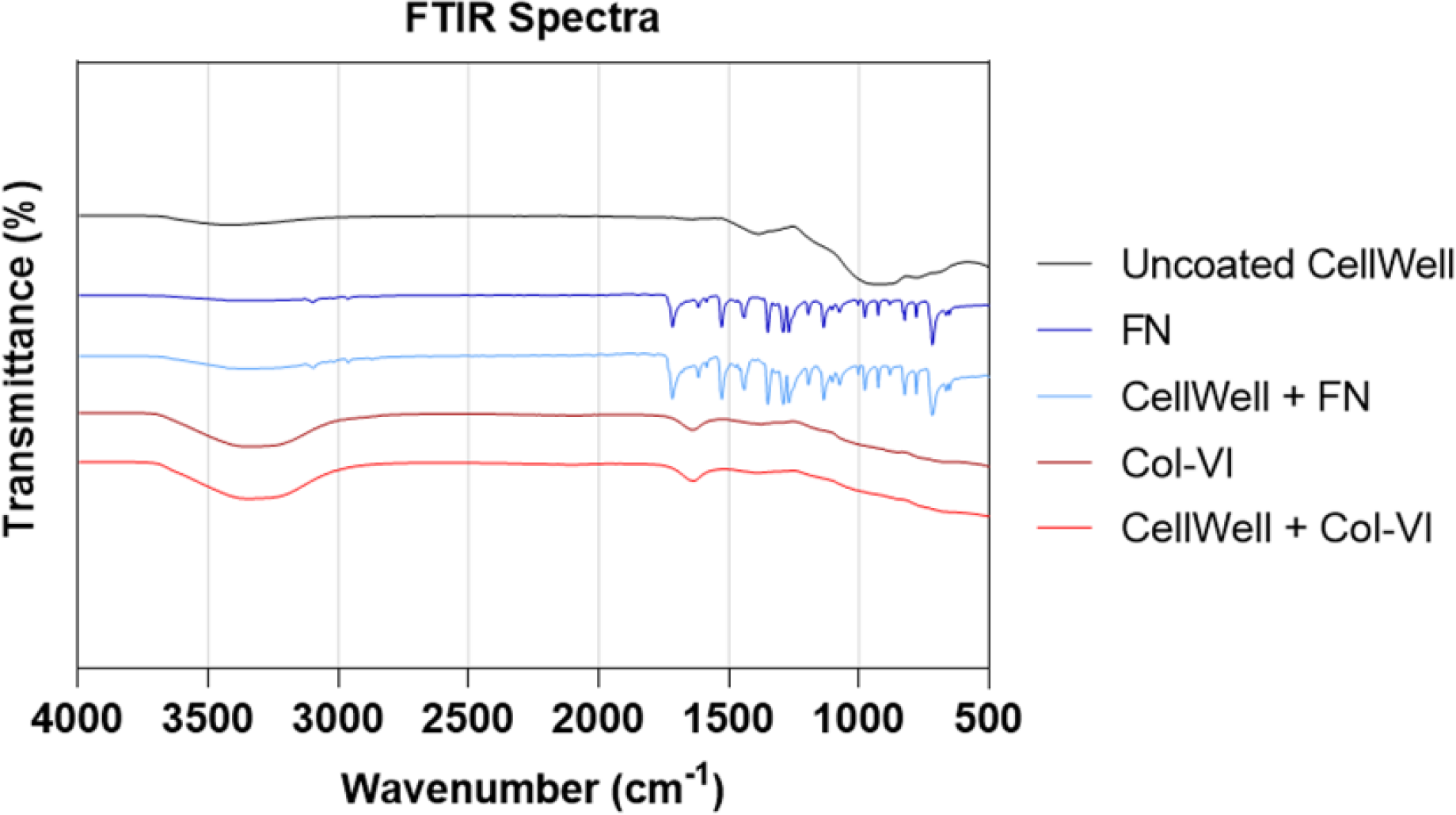
Successful coating of PCM proteins on the CellWells as confirmed by FTIR spectroscopy.

### 3.6 Nanofiber Characterization

Since the nanofibers were embedded into the Agarose CellWells to model collagen II fibers within articular cartilage, it was essential to obtain the distribution of their diameters. To the authors’ knowledge, the diameters of collagen II nanofibers in ankle cartilage have never been reported; thus, their measurement was necessary here to optimize the conditions for electrospinning CellWell PVA nanofibers. Fig. 9(A) and (B) show representative TEM images of both crosslinked PVA nanofibers and ankle articular cartilage, respectively. As seen in Fig. 9(C) the collagen II nanofibers had a median diameter of 50 nm compared to the 60 nm median diameter of PVA nanofibers. The PVA nanofibers were found to be within 10 nm for the median as well as the 25^th^ and 75^th^ quartiles of the ankle collagen II nanofibers as well, substantiating the use of PVA nanofibers to model the collagen II nanofibers in the CellWell.

**Fig. 9:**
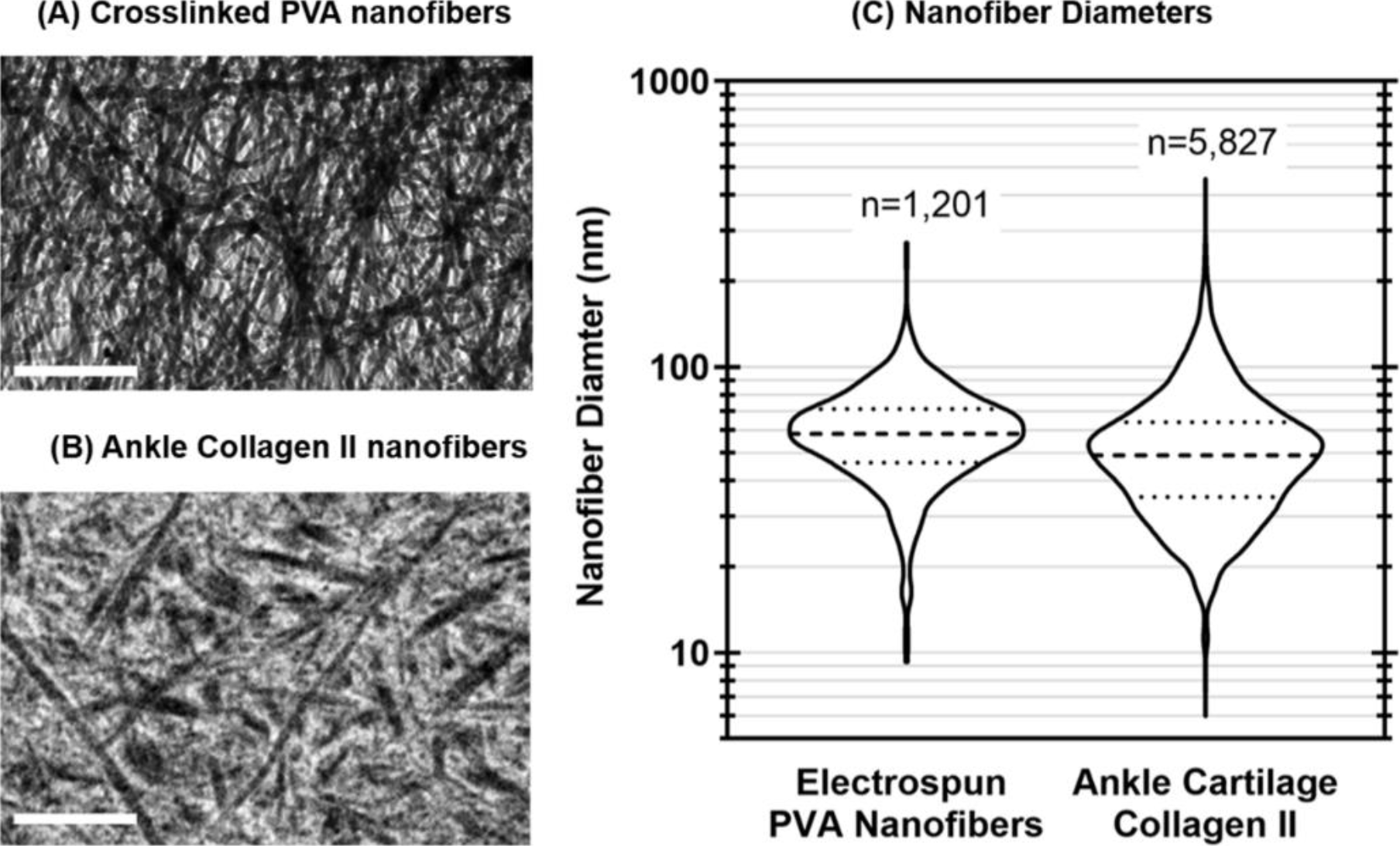
Electrospun PVA nanofibers have a diameter distribution representative of Collagen II fibers in ankle articular cartilage. TEM images of (A) PVA nanofibers post-crosslinking, and (B) collagen II fibers, extracted from the ankle; scale bars 1 μm. (C) The distribution of crosslinked PVA nanofibers to recapitulate nature of the collagen II fibers of the extracellular matrix of articular cartilage within our CellWell articular cartilage model for isolated chondrocyte cell culture. Fiber measurements were obtained from n=3 independent samples, with individual measurements represented on the graph.

### 3.7 Chondrocyte Viability

A fluorescent viability assay (ReadyProbes^®^ Cell Viability Imaging Kit, ThermoFisher) was conducted to assess both the cytotoxicity of the CellWell and its compatibility for use with standard live-cell imaging techniques. At 24-hours post-seeding, viability of 85.6% ± 10.5% (S.D.) was observed.

### 3.8 Chondrocyte Morphology

As depicted in Fig. 11, we have found that the CellWell is highly effective at promoting a physiological rounded chondrocyte morphology at 24 hrs of culture, as measured against standard 2D culture and 3D culture controls in which chondrocytes were embedded within agarose. Fig. 11 shows phase-contrast images and aspect ratios of chondrocytes seeded in agarose CellWells and control substrates – atop tissue culture polystyrene or agarose (2D culture) or encapsulated within agarose (3D culture). It can be easily seen that within 24 hrs post-seeding, many chondrocytes in 2D culture had lost their canonical spheroid morphology and started to spread. On the other hand, the chondrocytes in CellWells maintained their canonical morphology similar to those in 3D agarose culture. No statistical difference was observed between 3D agarose and CellWell chondrocyte morphologies, indicating maintenance of physiological morphology by the CellWell, while all other samples were found to be significantly different from each other.

## 4. Discussion

We have presented here a unique micropatterned nanocomposite cell culture platform to model articular cartilage that is suitable for high-throughput single-cell analyses of live cells using standard imaging techniques. All the images in Figs 10 and 11 were obtained using a standard inverted epifluorescence microscope. Contrary to some 2D micropatterned approaches [49], we do not have any non-adhesive areas to promote the cells being in a pattern. The geometry and spacing of our wells naturally promote the chondrocytes to fall into the wells.

**Fig. 10:**
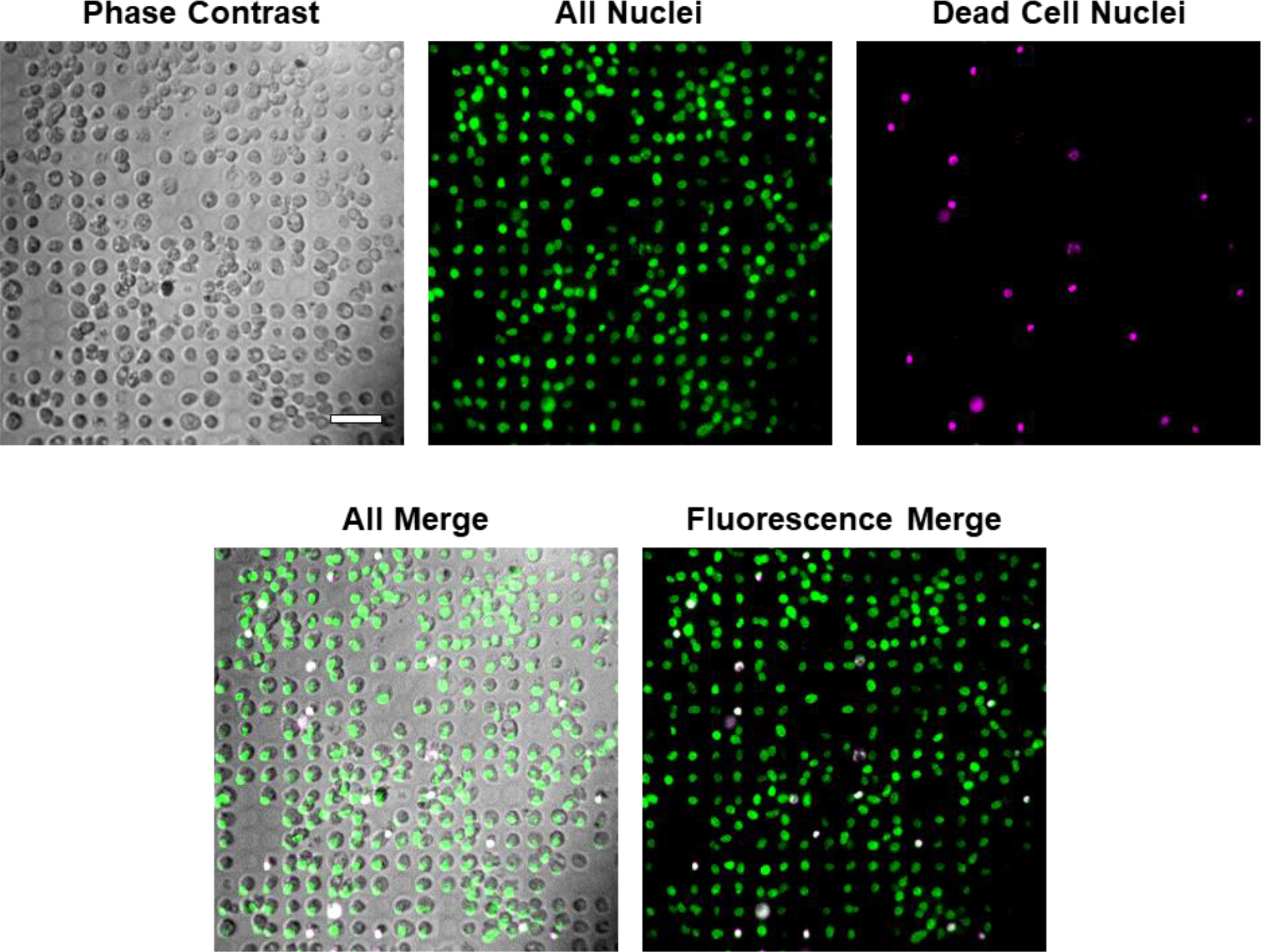
Chondrocyte viability is maintained in the CellWell. Chondrocyte viability of 85.6% ± 10.5% (S.D.) was observed at 24 hrs using a standard inverted epifluorescence microscope. Cell-permeable NucBlue stains all nuclei (shown in green), while cell-impermeable NucGreen stains only the nuclei of (dead) cells whose membranes have been disrupted (shown in magenta; dead cells appear white in overlaid images). Scale bar 50 μm.

**Fig. 11:**
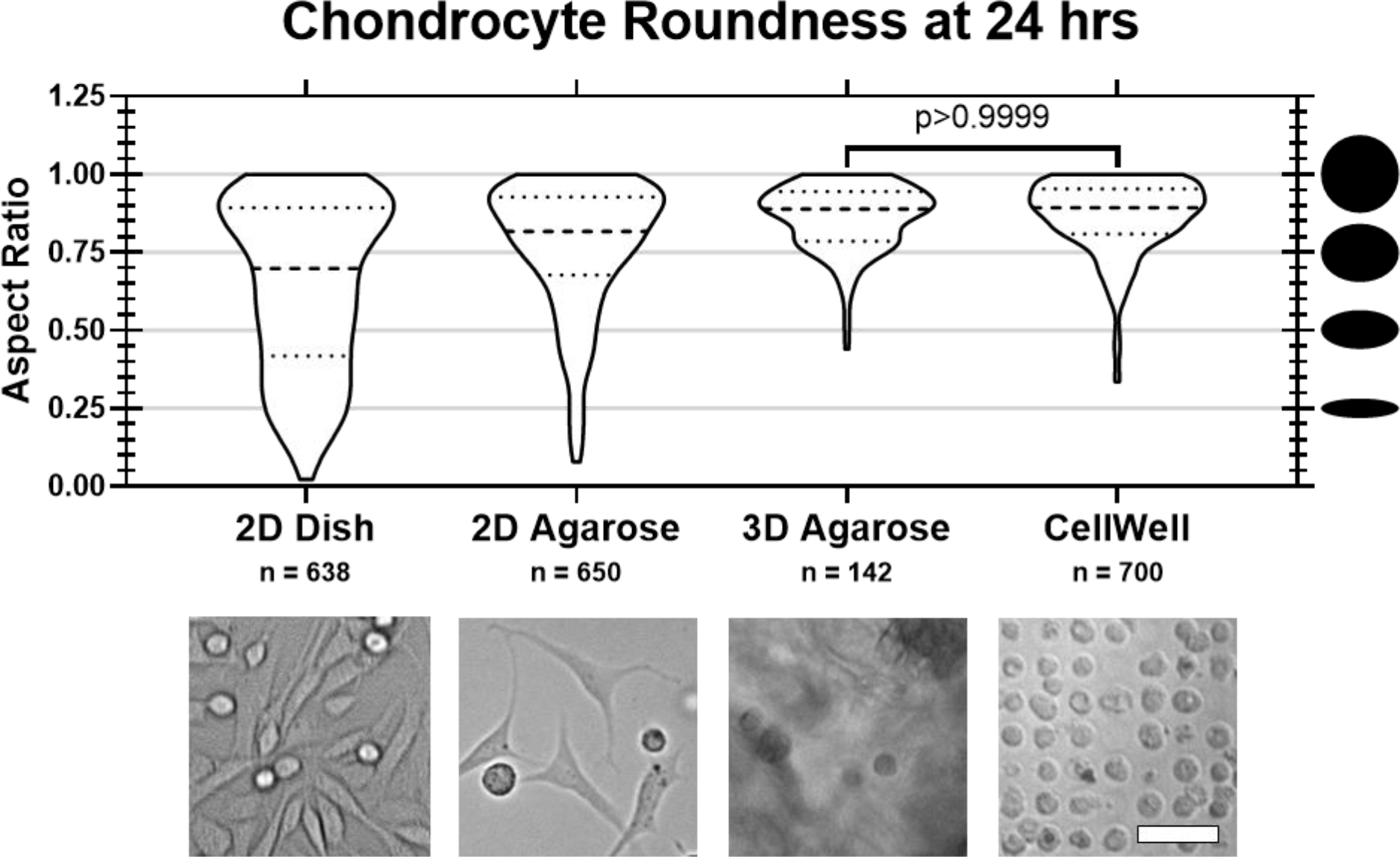
CellWells promote maintenance of physiological chondrocyte morphology. (A) Phase-contrast images of chondrocytes seeded using various platforms at 24 hrs are shown below distributions of aspect ratios of chondrocytes seeded with each platform from n=3 donors. No difference was observed between 3D agarose and CellWell chondrocyte morphologies, indicating CellWell maintenance of physiological morphology, while all other samples were significantly different from each other (p≤0.0002) based on Kruskal-Wallis with Dunn’s multiple comparisons post-hoc test. Scale bar 50 μm.

Two-dimensional substrates coated with non-adhesive polymers (e.g., poly[ethylene glycol] [PEG] or poly[2-hydroxyethyl methacrylate] [poly(HEMA)]), 2D substrates micropatterned with small circular islands of ECM proteins or amine groups surrounded by non-adhesive polymers, and suspension culture have all been shown promote maintenance of a rounded morphology and expression of phenotypic markers in chondrocytes, [30, 50, 51] however they do so at the cost of limiting adhesion. This lack of adhesion can have unintended and unpredictable consequences on cell signaling behavior – especially in studies of integrin-mediated mechanotransduction, which is well-known to be an essential mechanism of mechanotransduction in many cell types but has not been extensively examined in chondrocytes. Furthermore, this previous micropatterning technique relies on culturing cells directly on a coverslip, which adversely affects the mechanical environment of the chondrocytes since the coverglass stiffness is ~10^5^ Pa higher than that of the articular cartilage and ~10^6^ Pa higher than that of the chondrocyte PCM.

3D culture platforms (e.g., multicellular spheroids, organoids, and scaffolds [52] have also been shown to promote maintenance of chondrocyte phenotype [51, 53–60]; however, these techniques severely restrict the number of compatible analytical techniques, especially those capable of observing sensitive post-translational modifications of proteins that are key to cell signaling pathways. Moreover, 3D cultures are inherently difficult to image cells within, with high-quality images typically only attainable along the surface of samples (as in Fig. 1(A)).

To combine the advantages of 2D and 3D cell culture techniques, we have presented a unique micropatterned nanocomposite cell culture platform, the CellWell. The CellWell models the ECM of articular cartilage by utilizing a substrate composed of a hydrogel (to model cartilage proteoglycans) embedded with nanofibers (to model cartilage collagen II nanofibers). It is further micropatterned with a network of wells that are designed with geometries intended to reinforce the canonical spheroidal chondrocyte morphology. The CellWell addresses the problems of previous micropatterning techniques by providing a micropatterned environment that is much closer to the stiffness of the PCM than coverglass or tissue culture plastic and can promote adhesion.

One recent study utilized a similar philosophical approach to the CellWell by using ion milling of agarose to generate “cell hotels” for individual E. coli cells [61]. However, the agarose used in that study had a thickness of ~1 mm, which precluded its use for live-cell imaging. By utilizing a thin basal layer of only ~8 μm between the well bottoms and the underlying rigid substrate (e.g., coverglass or tissue culture dish), the CellWell provides a unique system that nestles only one cell per well, thus facilitating high throughput single-cell analysis of live cells with standard imaging techniques.

Another advantage of the CellWell over previous micropatterning techniques is the relatively minimal distance between wells. Instead of utilizing a large fixed distance of 30 μm or greater between any two consecutive wells [31, 44], the CellWell provides a range of distances between consecutive wells ranging from a minimal distance of 2.5 μm (slightly larger than the separation between cell pairs within chondrons *in vivo*) to a maximum distance of 15 μm. This, in combination with the fact that the wells are only approximately half the height of the cells, provides cells with the flexibility to either directly contact their neighboring cells by reaching over the space between wells or to remain in isolation. This flexibility provides immense possibilities for researchers interested in understanding the effects of cell-cell contact mechanics. Another advantage of the CellWell is the variable well diameters. Chondrocytes, like most cells, have a wide range of sizes. By providing a platform with a representative distribution of well diameters, the CellWell can provide a more natural environment for any given cell than systems with single fixed diameters.

In the current study, we have provided proof-of-principle for the CellWell as an articular cartilage model to prevent de-differentiation of articular chondrocytes that also facilitates high-throughput live-cell imaging studies. In future studies, we will seek to optimize the CellWell for long-term culture, allowing us to validate the phenotypic expression of chondrocyte marker proteins. We will also endeavor to develop a second-generation CellWell with multiple well geometries to model different depth zones of the articular cartilage.

## 5. Conclusions

We have demonstrated our ability to successfully synthesize a novel biphasic micropatterned platform – the CellWell. The compressive modulus of the CellWell was very close to that previously reported for the pericellular matrix of knee cartilage [44]. Ankle collagen II nanofiber diameters have been reported for the first time, and our crosslinked electrospun PVA nanofibers were found to have a diameter distribution representative of collagen II fibers. The micropatterned hemispheroidal wells in the CellWell promoted the canonical spheroidal morphology of articular chondrocytes with reasonable cell viability observed. Single-cell imaging was easily performed using a standard 20X objective on an inverted epifluorescence microscope due to the optical translucency of the CellWell (essentially a micropatterned thin film), and the embedded nanofibers were not found to inhibit optical translucency. These findings substantiate the applicability of the CellWell for use with standard culture and single-cell analysis techniques for live-cell imaging. The CellWell also has the advantage of material flexibility, with synthesis demonstrated using multiple hydrogel materials (i.e., agarose and PVA). Ultimately, we expect that by maintaining the physiological morphology of chondrocytes, the CellWell will promote physiological arrangement of intracellular structural proteins, thereby enhancing the clinical translatability of future studies conducted using this culture platform.

## Acknowledgments

We thank Dakota Lions Sight and Health, the Gift of Hope Tissue and Organ Donor Network, Dr. Susan Chubinskaya, Dr. Richard Loeser, and donor families for providing normal donor tissue. We thank Brian Baker at the Utah Nanofab for his technical assistance in providing the micropatterned silicon wafers. This material is based upon work supported by the National Science Foundation/EPSCoR Cooperative Agreement #IIA-1355423 and by the State of South Dakota. Any opinions, findings, and conclusions or recommendations expressed in this material are those of the author(s) and do not necessarily reflect the views of the National Science Foundation.

## Data Availability

The raw/processed data required to reproduce these findings cannot be shared at this time as the data also forms part of an ongoing study.

